# Intraspecific variation in symbiont density in an insect-microbe symbiosis

**DOI:** 10.1101/2020.11.03.365353

**Authors:** Benjamin J. Parker, Jan Hrček, Ailsa H.C. McLean, Jennifer A. Brisson, H. Charles J. Godfray

## Abstract

Many insects host vertically-transmitted microbes, which can confer benefits to their hosts but are costly to maintain and regulate. A key feature of these symbioses is variation: for example, symbiont density can vary among host and symbiont genotypes. However, the evolutionary forces maintaining this variation remain unclear. We studied variation in symbiont density using the pea aphid (*Acyrthosiphon pisum*) and the bacterium *Regiella insecticola*, a symbiont that can protect its host against fungal pathogens. We found that relative symbiont density varies both between two *Regiella* phylogenetic clades and among aphid ‘biotypes’. Higher-density symbiont infections are correlated with stronger survival costs, but variation in density has little effect on the protection *Regiella* provides against fungus. Instead, we found that in some aphid genotypes, a dramatic decline in symbiont density precedes the loss of a symbiont infection. Together, our data suggest that the optimal density of a symbiont infection is likely different from the perspective of aphid and microbial fitness. *Regiella* might prevent loss by maintaining high within-host densities, but hosts do not appear to benefit from higher symbiont numbers and may be advantaged by losing costly symbionts in certain environments. The standing variation in symbiont density observed in natural populations could therefore be maintained by antagonistic coevolutionary interactions between hosts and their symbiotic microbes.

## Introduction

Animals vary in the composition and abundance of the microbial communities that they harbor. Many microbial symbionts associated with insects, for example, are not found in all individuals within a species (Ferrari & Vavre, 2011). These facultative microbes can benefit their hosts (Brownlie & Johnson, 2009) but also impose metabolic costs and can require regulation through immunological control (Douglas, 2014; Herren & Lemaitre, 2011; Login et al., 2011; Pan et al., 2018). In light of these trade-offs, theoretical (Jaenike, 2012) and empirical studies (Hansen, Jeong, Paine, & Stouthamer, 2007; Jaenike, Unckless, Cockburn, Boelio, & Perlman, 2010) have considered how the prevalence of facultative symbionts across hosts changes in response to ecological factors. In addition, the mechanisms underlying these partnerships might evolve, for example leading to changes in the density of symbiont infections within hosts. For natural selection to shape host-symbiont associations there must be heritable variation in the traits affecting the partnership and this variation must be linked to host and/or symbiont fitness.

Recent studies have found that variation among microbial strains and host genotypes influences symbiont density and have identified specific molecular mechanisms controlling symbiont numbers. For example, different strains of the heritable bacteria *Wolbachia* transferred into a single *Drosophila simulans* line establish at different densities, and density is correlated with the protection conferred by *Wolbachia* against viruses (Martinez et al., 2017). Within the *Wolbachia* strain *w*MelPop, a genomic region termed Octomom was shown to influence density—artificial selection for higher Octomom copy number led to increased *Wolbachia* numbers in *Drosophila melanogaster* and to higher costs for its host (Chrostek & Teixeira, 2015). In the *Wolbachia*– *Drosophila* system, symbiont strain seems to be a more important factor than host genotype driving variation in within-host bacterial density, but other studies have emphasized the importance of host genetic variation in determining symbiont density. A gene called *Wolbachia density suppressor* (*Wds*) in the parasitic wasp *Nasonia* controls the density of *Wolbachia* through suppression of vertical transmission of the bacteria, and explains much of the variation in density between two closely related *Nasonia* species (Funkhouser-Jones, van Opstal, Sharma, & Bordenstein, 2018). This strain of *Wolbachia* is a clear reproductive parasite, and *Wds* was shown to be under positive selection. Another example comes from the pea aphid (*Acyrthosiphon pisum*) that we study here, where host genotypes vary in the density at which they maintain their obligate primary symbiont *Buchnera aphidicola* (Chong & Moran, 2016; Vogel & Moran, 2011; Wilkinson & Douglas, 1998).

What remains unclear, however, is what the existence of standing variation in symbiont density implies about the evolution of beneficial host-symbiont interactions. Is the optimal density of symbionts within a host the same from the perspective of the host and microbe? One possibility is that hosts and microbes have the same optimum symbiont density, but we find variation in symbiont density because of mismatches between host and symbiont genotypes—either in natural combinations or in those made artificially in the laboratory—that would be selected against in natural populations. Alternatively, hosts and symbionts might have different optimum densities, for example when higher symbiont density lowers host fitness but increases the probability of symbiont transmission. Variation in natural populations could then reflect an antagonistic process as hosts and symbionts struggle for control of symbiont density. To understand the evolutionary forces shaping a host-microbe association, we need to determine which partner is controlling the symbiosis and how variation among hosts and symbionts is linked with the fitness of both partners.

Here we study genetic variation in symbiont density using a tractable model host–microbe interaction: pea aphids and the facultative bacterial symbiont *Regiella insecticola*. In addition to their obligate symbiont *Buchnera*, aphids host a variety of bacteria that are found at intermediate frequencies in natural populations. These facultative symbionts are costly to maintain (Oliver, Campos, Moran, & Hunter, 2008; Vorburger & Gouskov, 2011) but also provide conditional benefits to their hosts. *Hamiltonella defensa*, for example, confers resistance against parasitoid wasps (Oliver, Russell, Moran, & Hunter, 2003) but decreases in frequency in the absence of infection pressure (Ives et al., 2020; Oliver et al., 2008). Similarly, *Regiella* makes aphids more resistant to natural fungal pathogens from the order Entomophthorales (Parker, Spragg, Altincicek, & Gerardo, 2013; Scarborough, Ferrari, & Godfray, 2005). Within pea aphids, *Regiella* forms two main phylogenetic clades (Henry et al., 2013), and the clades seem to have evolved to provide stronger protection against specific genotypes of the fungal pathogen *Pandora* in a genotype by genotype (G_Symbiont_ x G_Pathogen_) dependent manner (Parker, Hrcek, McLean, & Godfray, 2017).

The pea aphid species complex is comprised of multiple populations, termed ‘biotypes’, that are to some degree reproductively isolated (Peccoud et al., 2014; Peccoud, Ollivier, Plantegenest, & Simon, 2009) and are adapted to live on different species of plants in the Fabaceae family (Ferrari, Via, & Godfray, 2008; Via, 1991). Importantly, biotypes have been shown to differ in the frequencies at which they harbor facultative symbionts (Henry et al., 2013). Some biotypes are strongly associated with a particular symbiont species: for example, in collections from different continents *Regiella* is commonly found in aphids from *Trifolium spp*. (Ferrari, West, Via, & Godfray, 2012; Russell et al., 2013; Tsuchida, Koga, Shibao, Matsumoto, & Fukatsu, 2002). In contrast, some other biotypes rarely carry *Regiella*, or do not commonly harbor any facultative symbionts. To explain biotype-level patterns in symbiont frequencies, some studies have focused on the effects of symbionts on host-plant use with mixed success (McLean, van Asch, Ferrari, & Godfray, 2011; Tsuchida, Koga, & Fukatsu, 2004). Other studies have found that aphid and symbiont genotypes vary in the success with which they form artificial host-microbe pairings in the lab (Niepoth, Ellers, & Henry, 2018; Parker, McLean, Hrcek, Gerardo, & Godfray, 2017), suggesting that genotype may influence facultative symbiont frequencies in natural populations.

In this study, we tested a series of hypotheses about variation in relative density across both host and symbiont genotypes and the relationship between density and fitness. Our overall expectation was that *Regiella* density would be correlated with stronger protection against a fungal pathogen but higher survival costs to aphids. **(1)** We first tested the hypothesis that *Regiella* strains from two phylogenetic clades vary in within-host density, and **(2)** that higher density symbiont strains impose stronger survival costs on their hosts than do lower density strains. **(3)** We then tested whether the variation in density we uncovered among *Regiella* strains is dependent on host genotype (a G_Host_ x G_Symbiont_ interaction). This experiment used aphid lines that were naturally collected with a *Regiella* infection, and we next tested the hypothesis **(4)** that a single strain of *Regiella* would establish at different densities across a diverse panel of 28 host genotypes representing eight host-plant associated biotypes, some of which rarely harbor *Regiella*. **(5)** We then measured the effects of host genotype on the strength of protection conferred by *Regiella* against fungus and on aphid survival and **(6)** if these costs and/or benefits are correlated with symbiont density. We observed that many lines lost their *Regiella* infections during our experiment, and **(7)** tested the hypothesis that low density lines are in the process of losing their symbiont infections. We found a high degree of variation in density across both hosts and microbes. However, we did not find strong evidence that density is associated with the fitness effects of harboring *Regiella*. We instead found that symbiont loss is preceded by a drop in density and varies among genotypes. We discuss these findings in the context of host-symbiont coevolution.

## Methods

Pea aphids are cyclically parthenogenetic allowing us to maintain asexual genotypes in the laboratory under a light and temperature regime of 16L:8D and 14°C. Aphids were housed in Petri dishes with a *Vicia faba* leaf (cultivar ‘The Sutton’), a ‘universal’ host plant on which all aphid biotypes are able to feed (Sandström & Pettersson, 1994), with petioles inserted in 2% tap water agar. When new lines were collected from the wild, we screened them for seven pea aphid facultative symbionts using PCR and symbiont-specific primers (94°C 2m, 12 cycles x (94°C 30s, 56°C (declining 1°C each cycle) 50s, 72°C 90s), 26 cycles x (94°C 30s, 45°C 50s, 72°C 90s) and a final extension of 72°C 5m) (Henry et al., 2013). We cleared lines of any facultative symbionts by immersing the petioles of *V. faba* leaves in an antibiotic solution (100 mg ml^-1^ ampicillin, 50 mg ml^-1^ cefotaxime and 50 mg ml^-1^ gentomicin), and feeding first-instar aphids on the leaves for 48 hours (McLean et al., 2011); note that lines collected with facultative symbionts that we are currently unable to cure (*Spiroplasma* & *Rickettsia*) were not included in our experiments. After antibiotic exposure, we moved aphids to fresh leaves until they became adults. We collected the late offspring (>10 in birth order) of antibiotic-fed aphids, maintained them until they became adults, tested them for symbionts using PCR as above, and retained the offspring of symbiont-free individuals. We kept these lines for at least 15 generations to ensure there were no maternal effects of antibiotic treatment on experimental aphids, while periodically re-testing to ensure the lines were free of secondary symbionts.

Facultative symbionts can be introduced into aphid lines via microinjection, which allows us to create different combinations of host and symbiont genotypes. We established symbionts in new host lines by injecting a small volume of hemolymph (approximately 0.25μl) from an infected adult donor aphid into a one-day-old 1^st^ instar recipient using a capillary needle (Lukasik et al., 2015). Injected aphids were then reared on leaves of *V. faba* in Petri dishes at 14°C until they became adults when each aphid was moved into its own dish. These adults were allowed to reproduce, and approximately the 10^th^ offspring was reared to maturity, allowed to reproduce, and then frozen at -20°C. DNA extraction and PCR were used as described above to screen the lines for symbiont establishment. Lines that successfully acquired a *Regiella* infection were then reared individually and re-tested for infection before use in experiments.

### (1) Density of *Regiella* strains in a common host genotype

We tested the hypothesis that different *Regiella* strains in genetically identical hosts vary in their relative densities. As part of a different project, symbiont strains were collected from the United Kingdom and the United States, and six housekeeping genes were sequenced to build a phylogeny (Figure 1A) (Parker, Hrcek, et al., 2017). The 13 *Regiella* strains included representatives of the two major phylogenetic clades found in pea aphids (which separated ∼500,000 years ago (Henry et al., 2013)). In this previous study, we transferred each *Regiella* strain into an uninfected aphid recipient (genotype 145) using the protocol described above.

**Figure 1:**
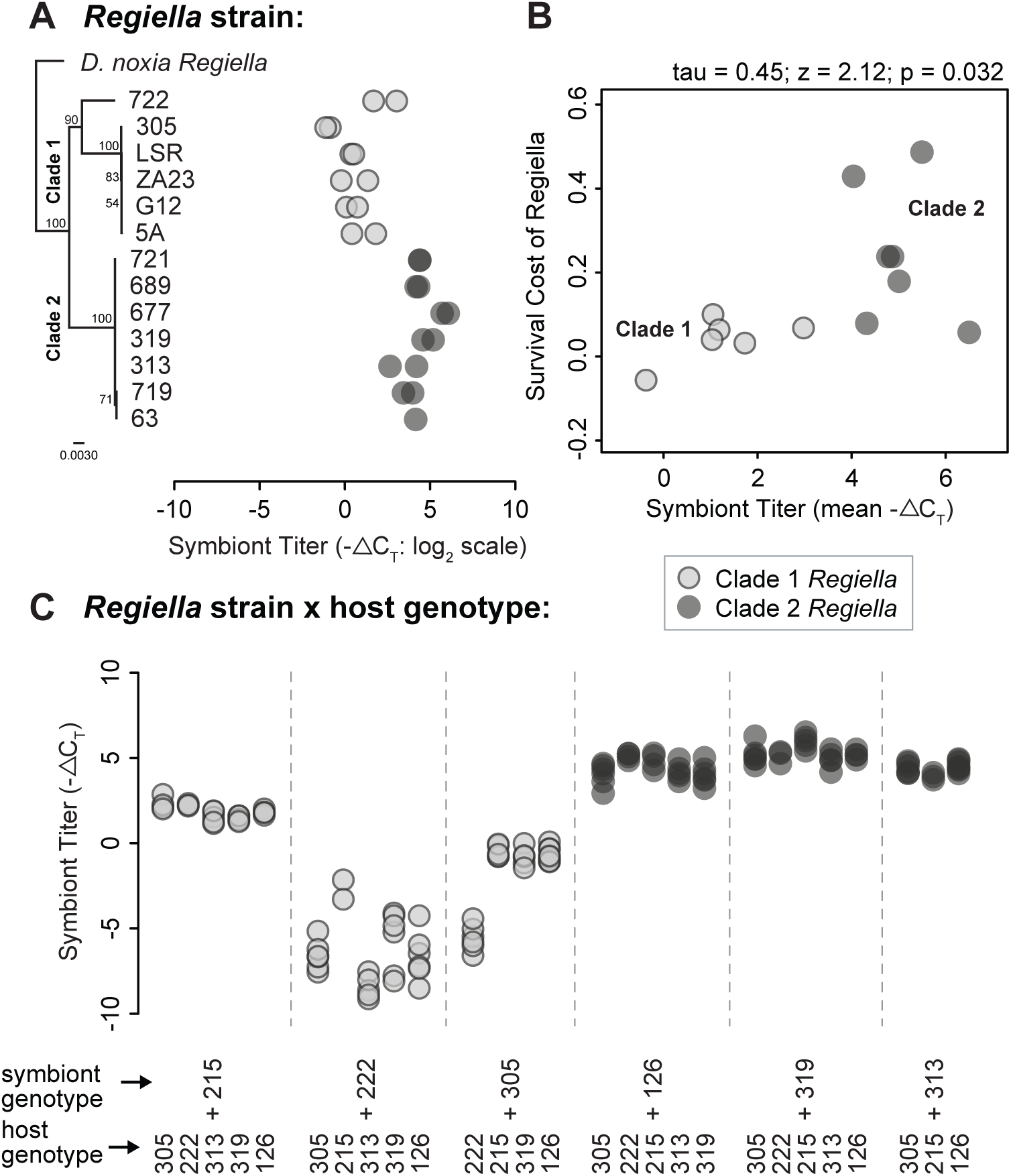
*Regiella* strains vary in the density at which they establish in hosts. **A:** The phylogeny of *Regiella* strains is shown along the Y-axis, along with the strain names. A *Regiella* strain from another aphid species (*Diuraphis noxia*) was used to root the phylogenetic tree. -ΔC_T_ (critical threshold) values measured using qPCR are shown along the X-axis, which can be interpreted as the relative symbiont densities of each sample on a log_2_ scale. Each of the biological replicates (a DNA extraction from a single adult aphid) for each strain are shown as points on the plot. Clade 1 and 2 genotypes shown in light and dark grey points, respectively. **B:** This plot shows the relationship between symbiont density (as shown by -ΔC_T,_ on the X-axis) and survival cost (fraction of aphid without *Regiella* surviving the experiment - fraction with *Regiella* surviving on the Y-axis). The mean value for each genotype is shown by a single point, with the Clade 1 and 2 genotypes shown in light and dark grey, respectively. **C:** This graph shows the density of multiple *Regiella* strains established in different host genotypes. The Y-axis shows symbiont density as above, with each point representing a biological replicate. Symbiont and host genotype names are shown along the bottom of the figure as indicated.

In this study, we measured relative symbiont density using quantitative PCR (qPCR). We reared lines for eight generations and froze adult aphids (11 days old) individually from each line at -80°C. We extracted DNA from each aphid using the Qiagen DNEasy kit under recommended conditions. We designed qPCR primers based on a conserved region of the *hrpA* gene (F: CGCATTGGGAGAAAAGCCAAG; R: CCTTCCACCAAGCCATGACG). We compared amplification of *Regiella hrpA* to Glyceraldehyde-3-phosphatedehydrogenase (*g3PDH*) in the aphid genome (F: CGGGAATTTCATTGAACGAC; R: TCCACAACACGGTTGGAGTA). It is important to note, therefore, that our measurements of symbiont density are relative to the number of host cells in a sample. PCR conditions for both sets of primers were optimized against a 1:10 serial dilution of gDNA (200ng – 0.2ng gDNA per reaction) to 99.2% for *hrpA* and 100.0% for *g3PDH*, and were ultimately run at concentrations of 400nM F/350nM R and 300nM for *g3PDH* and *hrpA*, respectively. qPCR reactions were run on a Bio-RAD CFX96 Real-Time System machine, with an initial step of 95°C for 3 minutes and 40 cycles of 95°C for 10s and 60°C for 30s. Each 20μL reaction included a 1X PCR buffer, Mg^+2^ at 2mM, dNTPs at 0.2mM, EvaGreen at 1X, 0.025 units/μL of Invitrogen taq, and 40ng of gDNA. We ran three technical replicates for each reaction.

We calculated a -ΔC_T_ value for each sample by subtracting the mean C_T_ value of the three technical replicates for *g3PDH* from the mean C_T_ value of *hrpA*. We analyzed -ΔC_T_ values using a 2-factor ANOVA after testing for model assumptions using R v.3.4.1. *Regiella* strain was nested within clade, and we analyzed the main effects of clade and strain.

### (2) Correlation between *Regiella* density and aphid survival

In a previous study, we quantified the survival costs of carrying *Regiella* in these same 13 strains in a single aphid host genetic background (Parker, Hrcek, et al., 2017). Briefly, we calculated the percentage of aphids harboring each strain of *Regiella* that survived over an 8-day period, and we subtracted this value from the survival of symbiont-free aphids of the same genotype to calculate survival cost. Here, we used this survival data together with the symbiont density data to test the hypothesis that higher density symbiont strains impose stronger costs on their hosts. We calculated a non-parametric (Kendall’s rank-order) correlation coefficient to look for an association between symbiont density and survival cost using the statistical package R v.3.4.1. We first analyzed all of the strains together in a single analysis, and then repeated the analysis for the strains in the two major clades separately.

### (3) GxG effects on *Regiella* density

Using a second panel of lines generated previously (Parker, Hrcek, et al., 2017), we tested the hypothesis that variation in density among *Regiella* strains varies across host genotypes (a G_Host_ x G_Symbiont_ interaction). Each of the six aphid genotypes included in this panel had a ‘natural’ *Regiella* infection when originally collected, and the aphid lines belonged to two biotypes (associated with the host plants *Trifolium pretense* or *Medicago sativa*). Six *Regiella* strains were established in each of the six host genotypes as described above (except for three combinations which failed). We focused our study of symbiont density only on the novel pairings of host and symbiont genotypes (which were generated using injection) and did not include the ‘natural’ pairings. Samples were collected and *Regiella* densities were measured using qPCR as above. We analyzed the -ΔC_T_ values using a generalized linear model with a quasi-likelihood distribution in R v.3.4.1. The model included terms for *Regiella* genotype, host genotype, and the interaction between these terms. We performed model comparisons using ANOVA, by first removing the interaction term, then host genotype, then symbiont genotype.

### (4) *Regiella* density across diverse aphid biotypes

We found a significant but small effect of host genotype on *Regiella* density in the experiment above; we then tested the hypothesis that the density of a single strain of *Regiella* would vary across a more diverse panel of genotypes from multiple host-plant associated biotypes, some of which rarely harbor *Regiella*. We collected genotypes from 8 different host plants from field sites in southern England (Table S1). Aphids were assigned to biotypes using microsatellite sequencing (Hrcek et al., 2018). Lines were tested for symbionts and these were then removed as described above. We then established a single strain of *Regiella* (strain 313) in each aphid genotype using hemolymph transfers as above, and we confirmed successful *Regiella* transfer by PCR detection. We reared aphids in the lab for eight generations, at which point we screened the lines for *Regiella* again. This procedure often required multiple replicate injected lines of an aphid genotype because it was difficult to establish *Regiella* in many of the lines and some replicates lost their symbiont infections by generation eight. This protocol was successful for 28 aphid genotypes, from which we then collected three adult aphids each, extracted DNA, and measured *Regiella* density using qPCR as above.

### (5) Host genotype effects on fungal protection and survival costs

Using the panel of 28 aphid lines with and without *Regiella*, we tested the hypothesis that the strength of symbiont mediated protection against *Pandora* and the survival costs of harboring *Regiella* varied across host genotypes. We obtained strains of the fungal pathogen *Pandora neoaphidis* from the USDA Agricultural Research Service’s Collection of Entomopathogenic Fungal Cultures. We maintained fungal cultures in the laboratory by infecting aphids from a symbiont-free aphid clone that is a genomic mixture of multiple aphid biotypes every two weeks (genotype 145), and we stored dried cadavers at 4°C. Fungal cadavers were hydrated to induce sporulation by placing them on tap water agar for 12h overnight. We used spores from *Pandora* strain ARSEF2588 for the main fungal infection.

We infected aphids from the 28 host genotypes with and without *Regiella* strain 313 in a single temporal block. Aphids were reared on *V. faba* plants in cup-cages. We subjected adult aphids (11 days old) to a ‘spore shower’ from two fungal cadavers in an infection chamber. Fungal cadavers were rotated among the infection chambers so that each cadaver was present for the same amount of time above each chamber to ensure equal spore dose and infection risk across aphids. We have used this protocol previously to control for fungal cadaver age, pathogen dose, and aphid cadaver quality (Hrcek et al., 2018; McLean et al., 2018; Parker, Hrcek, et al., 2017). Control aphids were handled similarly but were not exposed to fungal spores. Between 23 and 106 aphids (mean 90.6) were used per line depending on availability. After infection, we maintained aphids in dishes as above in groups of five. Each dish was wrapped in parafilm to keep the humidity high (>95%) and kept at 20°C for 48 hours. We then moved aphids to new dishes without parafilm and with fresh plant material. The experiment was blinded (i.e. observers were unaware of aphid genotype, symbiont status or fungal exposure treatment), and pathogen infection frequencies and survival were recorded on day 7 after infection.

We analyzed sporulation status (whether an aphid produced visible signs of sporulation) using binomial-error, linear mixed-effects models implemented in the lme4 package in R v.3.4.1 (Bates, Mächler, Bolker, & Walker, 2015). Symbiont status (no secondary symbionts or *Regiella* strain 313) and aphid biotype were treated as fixed effects, with aphid genotype nested within biotype and modeled as a random effect. Minimal models were derived by first removing the interaction term followed by biotype and then symbiont, with the statistical importance of each term assessed using ANOVA. Post-hoc comparisons of aphid biotypes were made using Tukey’s HSD pair-wise comparisons in the multcomp package (Hothorn, Bretz, & Westfall, 2008) in R. We analyzed aphid survival to assess the costs of harboring *Regiella* using the same methods.

We repeated this experiment using six host genotypes, again with and without *Regiella* strain 313, this time using a different genotype of fungal pathogen (ARSEF6631). Sporulation status was recorded as above on day seven after infection as above. These data were modeled using binomial-error linear models (with aphid genotype and symbiont status as fixed effects) and Tukey’s HSD pair-wise comparisons.

### (6) Correlations between *Regiella* density and host fitness across aphid biotypes

We then tested the hypothesis that variation in density across diverse aphid genotypes is correlated with the protective benefits and survival costs of harboring *Regiella*. We calculated Kendall’s rank correlation coefficients (in R v.3.4.1) between symbiont density and fitness effects for the 28 genotypes in the fungal infection experiment. -ΔC_T_ values were used as the measurement of density as above. The strength of protection was calculated by (percent sporulated aphids with *Regiella*) / (percent sporulated aphids without *Regiella*) for each genotype. Survival costs were calculated with (percent survival with *Regiella*) / (percent survival without *Regiella*), with larger values indicating higher survival costs.

### (7) *Regiella* density and loss across aphid generations

In the experiments above we found several aphid lines with very low *Regiella* densities, and we noticed that many aphid lines lost their *Regiella* infections over the eight generations after symbiont establishment. We tested the hypothesis that these low-density lines are in the process of losing their symbiont infections by tracking symbiont density over multiple generations after injection. We selected two aphid genotypes from the previous experiments, one from the *Trifolium* biotype (genotype 317) and one from *Lotus corniculatus* (genotype 663). We selected these lines for further examination because in a previous study we found that aphids from *Trifolium* retain their symbionts at a higher frequency than do aphids from *Lotus corniculatus*. In the experiment above, genotype 317 from *Trifolium* had a much higher *Regiella* density (-ΔC_T_ = 2.86) than genotype 663 from *Lotus corniculatus* (-ΔC_T_ = -6.02) after eight generations, and we expected genotype 663 to have a low density of *Regiella* because it was in the process of losing its symbiont. We conducted the experiment in two blocks that were several months apart. We injected replicates of both genotypes (genotype 663: block 1, n = 9; block 2, n = 10; genotype 317: block 1, n = 3; block 2, n = 4) with *Regiella* strain 313 using the injection protocols described above. We then screened adult aphids from each replicate at the first generation after injection (generation 1) and each of the next four generations (generations 2-5). We collected 2-5 adult aphids from each line on the first day of reproduction at each generation and stored the samples at -20°C. We extracted DNA and screened each sample for *Regiella* using PCR protocols as described above. For samples with a positive PCR result indicating the presence of *Regiella*, we used qPCR to measure symbiont density at generation 2 and at generation 4. We compared the ratio of symbionts lost and retained between the two genotypes with a 2 x 2 contingency table and a Yates corrected Chi-squared test.

## Results

### (1) *Regiella* strains vary in density in a common host background

Using the panel of 13 *Regiella* genotypes in a common host background, we found that two *Regiella* clades significantly differed in density (ANOVA; clade: F = 212, 1DF, p < 0.0001; Figure 1A), and *Regiella* strains significantly varied in density independently from the effect of clade (Strain: F = 4.31, 11DF, p = 0.01; Figure 1A). Densities of Clade 2 *Regiella* were on average 13.3 times higher than those in Clade 1, and phylogenetic clade explained a much larger portion of the variance in our model (78.1% total variance) than did strain (17.5%).

### (2) Higher density is correlated with lower host survival

We found that higher *Regiella* densities are correlated with higher survival costs to hosts (tau = 0.45; z = 2.12, p = 0.032; Figure 1B). An important caveat, however, is that differences between the two clades other than density might impact host survival, and we did not find significant correlations between *Regiella* densities and survival cost among strains within each clade (Clade 1: tau = 0.33, z = 0.94, p = 0.35; Clade 2: tau = 0.20, z = 0.61, p = 0.54).

### (3) Variation in density due to symbiont genotype is not strongly dependent on host genotype (a G_Host_ x G_Symbiont_ interaction)

Using the panel of six *Regiella* genotypes in six host genotypes, we again found that Clade 1 *Regiella* established at lower densities than Clade 2 symbionts (Figure 1C). In our model, symbiont genotype (F = 1084, 5DF, p < 0.0001), host genotype (F = 57.0, 5DF, p < 0.0001), and their interaction (F = 15.4, 16DF, p < 0.0001) had statistically significant effects on density. Symbiont genotype explained 93.6% of the deviance accounted for by the full statistical model, while host genotype (2.1%) and the host genotype by symbiont genotype interaction (4.3%) explained less of the deviance in the model.

### (4) Aphid genotypes from diverse biotypes vary in *Regiella* density

Using 28 different host genotypes in which we established *Regiella* (strain 313), we found extensive variation in symbiont density due to host genotype (Kruskal-Wallis χ^2^ = 71.7, DF = 26, p < 0.0001; Figure 2A) and among aphid biotypes (Kruskal-Wallis χ^2^ = 26.8, DF = 7, p < 0.001; Figure 2A). Some lines were found to have very low *Regiella* densities that were several orders of magnitude lower than average (Figure 2A).

**Figure 2:**
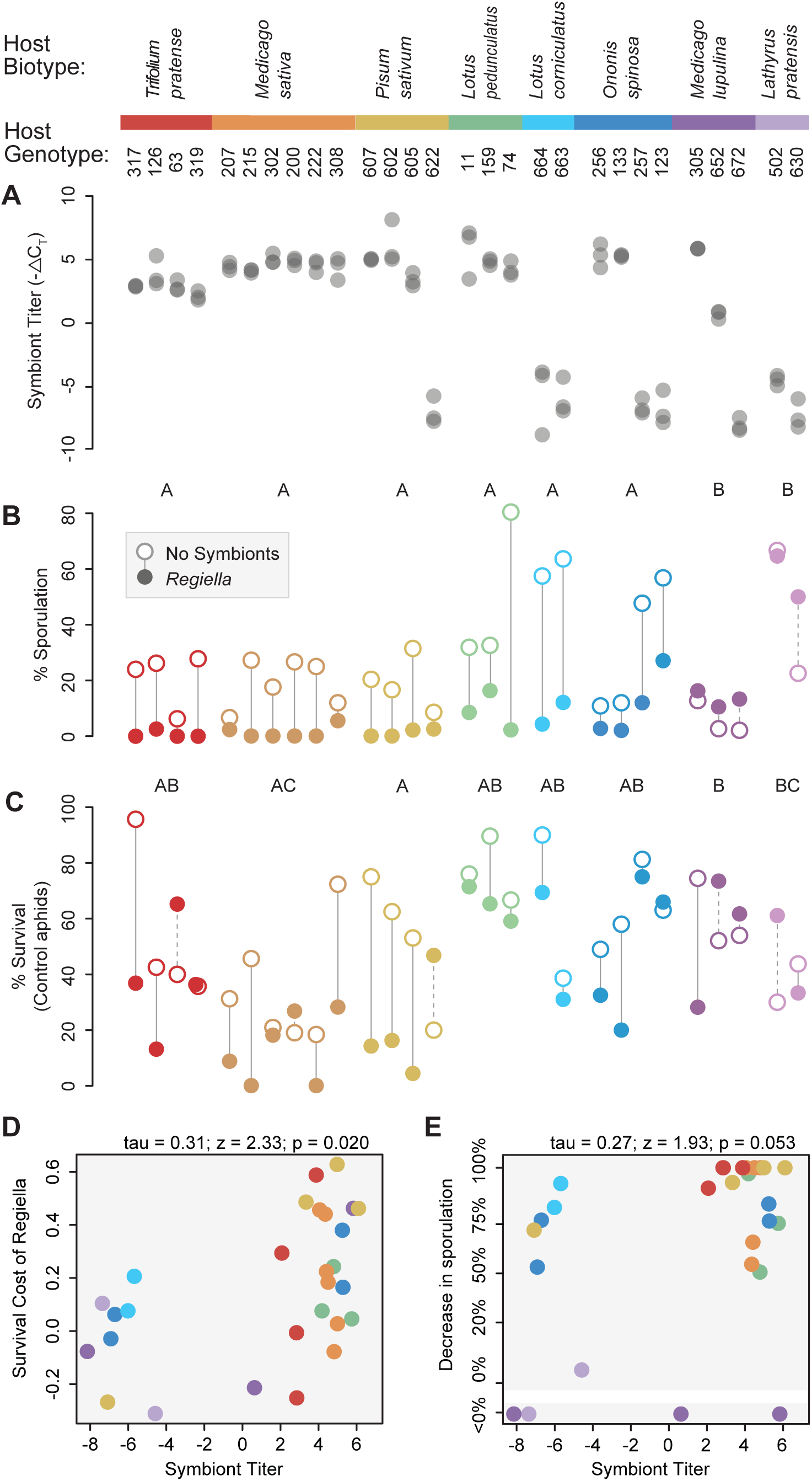
Host genotypes influence *Regiella* density and loss. The host biotype and genotype names for each line in used in these experiments is shown along the top of the figure. **A:** The Y-axis represents *Regiella* strain 313 density as -ΔC_T_ values, which can be interpreted as the relative symbiont density on a log_2_ scale. Each replicate is shown by a grey point. **B:** This plot shows sporulation data (the appearance of a sporulating fungal cadaver after 7 days) of aphid genotypes with (filled circles) and without (empty circles) *Regiella*. The strength of symbiont-mediated protection is reflected by the relative sporulation of lines with and without *Regiella* and is represented by a vertical grey line. Post-hoc tests comparing these differences across biotypes (interaction terms between biotype and *Regiella* presence) are shown along the top of the figure. **C:** The Y-axis shows the percent survival data of control aphids (not exposed to fungus) after 7 days. Circles and lines are as above. **D:** This plot shows the relationship between symbiont density and survival costs (percent survival without *Regiella* – percent survival with *Regiella*). **E:** This plot shows the relationship between symbiont density and protective benefit (percent sporulated aphids with *Regiella* / percent sporulated aphids without *Regiella*).

### (5) Host genotype influences the protective benefits and survival costs of harboring *Regiella*

Using the host genotype panel, we found that *Regiella* protected aphids from *Pandora* infection as expected (Figure 2B; Symbiont: χ^2^ = 169.32, 1DF, p < 0.0001). However, biotypes differed significantly in the extent of this protection (Symbiont * Biotype: χ^2^ = 89.9, 7DF, p < 0.0001; post-hoc comparisons Figure 2B; Table S2). Specifically, we found that harboring *Regiella* provided no protection against fungal infection in aphids from the *Medicago lupulina* and *Lathyrus pratensis* biotypes (post-hoc comparisons, Figure 2B; Table S2). A replicate experiment using a second fungal isolate (ARSEF6631) mirrored these results: protection against fungus was again strongly dependent on aphid host genotype, with the two genotypes from *Medicago lupulina* experiencing no protection from *Regiella* (Figure S1; Table S2). Host genotypes also differed in the severity with which they experienced survival costs due to *Regiella*, with this pattern differing significantly across aphid biotypes (Biotype x Symbiont: χ^2^ = 26.3, 7DF, p < 0.001; Figure 2C; post-hoc comparisons, Table S2).

### (6) The costs but not the benefits of symbiosis are correlated with density

Across the host biotype panel, we found that lines with higher *Regiella* densities experienced greater fitness costs of harboring *Regiella* (stronger decrease in survival of control aphids when harboring *Regiella*; tau = 0.31; z = 2.33; p = 0.020, Figure 2D), but the correlation between density and protection against fungal pathogens was not statistically significant (tau = 0.27; z = 1.93; p = 0.053, Figure 2E).

### (7) Symbiont loss is preceded by a drop in density and varies among genotypes

We observed that some aphid lines lost their *Regiella* infections after injection, and in a previous study we found that aphid genotypes varied in their ability to maintain *Regiella* across generations (Parker, McLean, et al., 2017). We hypothesized that the very low densities we measured in some aphid lines in the experiment described above could reflect lines that are in the process of losing their symbiont infections. In the experiment above, genotype 317 from *Trifolium* had a much higher *Regiella* density (-ΔC_T_ = 2.86) than genotype 663 from *Lotus corniculatus* (-ΔC_T_ = -6.02) after 8 generations. In this experiment, we found that no lines from genotype 317 (0 out of 7) lost their *Regiella* infection over 5 generations. Among lines from genotype 663, 11 out of 19 lines lost their symbiont. In 10 lines from genotype 663, loss was preceded by a sharp drop in symbiont density (Figure 3). The rate of symbiont loss differed significantly between the two genotypes (χ^2^ with Yates’ correction = 4.37, 1DF, p = 0.037).

**Figure 3:**
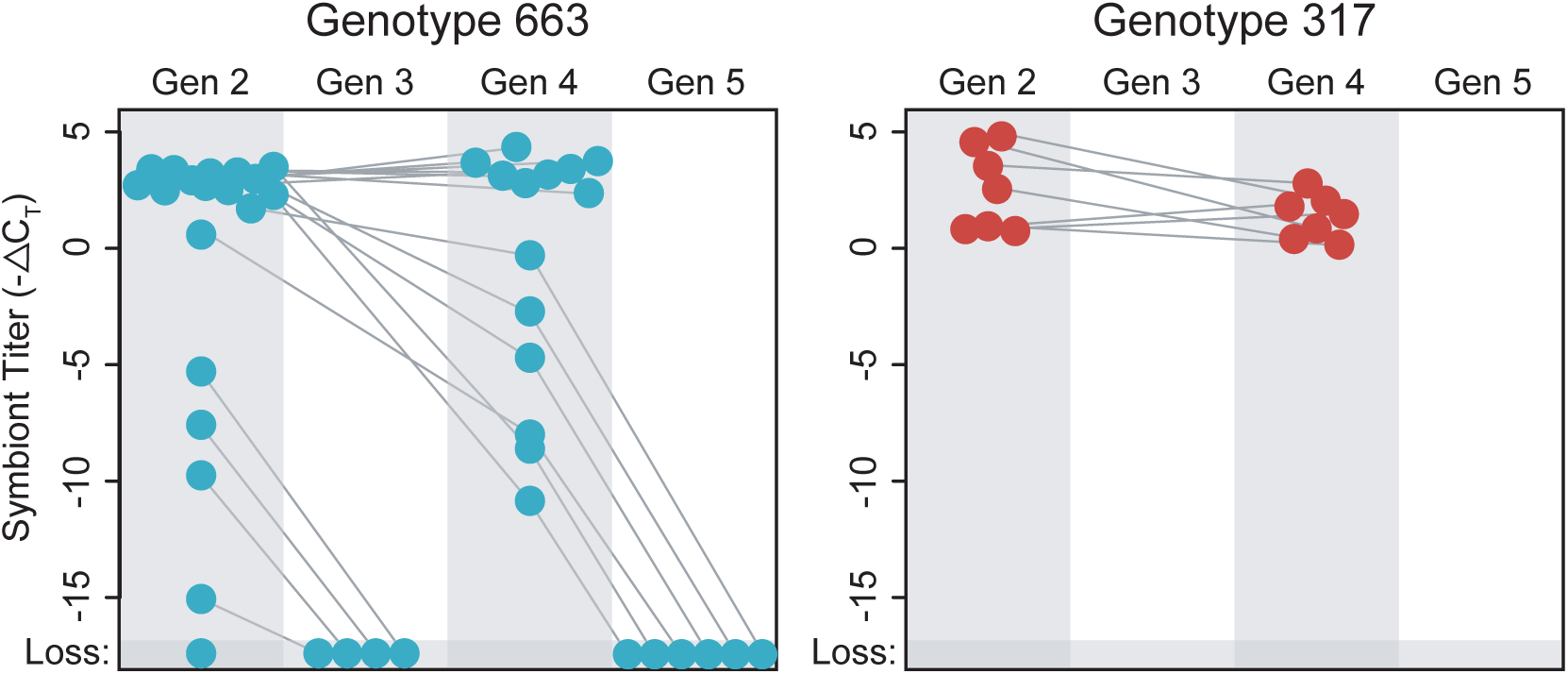
Symbiont loss over several generations. The y-axes show symbiont density as above, with samples taken at multiple generations shown along the top of the figure. Each point represents a different aphid line established with a *Regiella* infection from host genotype 663 (blue) or 317 (red) connected by a grey line. Lines that lost their symbiont infection as determined by PCR are shown along the bottom of the figure.

## Discussion

We confirmed that symbiont density varies extensively across both host and microbial genotypes. Strains from the two major *Regiella* clades, when introduced into a single host genetic background, were found to be present at average densities that varied by ∼13X (with clade explaining a large part of this variation). A similar pattern was found when *Regiella* strains from the two clades were introduced into multiple host genotypes. *Regiella* densities were quite similar across those aphid genotype which when collected in the field naturally harbored this symbiont (prior to being cleared before the experiments). But we found much more extensive variation in symbiont density across aphid genotypes from multiple biotypes that had not all originally carried *Regiella*. We found that low densities of symbionts are frequently a prelude to the complete loss of the association, and that the likelihood this occurs is influenced by host genotype.

Across host genotypes, the strength of symbiont-mediated resistance to *Pandora* was not significantly correlated with *Regiella* density: some aphid lines with a high symbiont density experienced no protection against *Pandora* infection, while other lines with fewer symbionts were well-protected. Variation in density across *Regiella* strains may not be correlated with fungal protection because, in this system, there is a symbiont strain by fungal genotype (GxG) interaction (Parker, Hrcek, et al., 2017) with particular *Regiella* strains conferring strong protection only against specific *Pandora* genotypes. Other studies have found correlations between symbiont density and protection from pathogens across symbiont genotypes (e.g. Chrostek et al., 2013; Martinez et al., 2015; Martinez et al., 2017). A better understanding of when density influences symbiont-mediated protection will be aided by more examples of the molecular mechanisms underlying protection (Gerardo & Parker, 2014). One possibility is that some types of mechanisms, like the production of an anti-pathogen toxins (e.g. (Ballinger & Perlman, 2017; Oliver, Degnan, Hunter, & Moran, 2009)), are density-dependent while others, like protection that is mediated by the host’s immune system (e.g. Hughes, Koga, Xue, Fukatsu, & Rasgon, 2011; Kambris et al., 2010), are not influenced by symbiont density.

Evidence for our hypothesis that symbiont density is correlated with fitness costs to hosts was mixed. Survival costs were significantly correlated with *Regiella* density across host genotypes, but there were a number of lines with a high density *Regiella* infection that experienced no measurable costs of symbiont infection. Across *Regiella* strains, higher density infections were correlated with higher survival costs to aphids. But other genetic differences between the two *Regiella* clades might be driving both costs to hosts and differences in density, and we found no correlation between density and survival within the symbiont clades. These results might suggest that *Regiella* from the two clades, which are separated by a half million years (Henry et al., 2013), have adopted different strategies for surviving within hosts, i.e. Clade 2 *Regiella* establish at higher densities in hosts but are more costly to hosts. It is possible that higher density *Regiella* infections might protect symbionts against failure to be transmitted to offspring or increase the probability of horizontal transmission (Henry et al., 2013), but come at the expense of host survival.

By following infections across multiple generations, we found that low density infections tend to precede symbiont loss, and that the rate of low-density followed by symbiont loss varies between two genotypes. Aphid facultative symbionts are thought to be transmitted with high fidelity from mothers to offspring, but recent studies have shown how host biotype (Parker, McLean, et al.) and donor and recipient relatedness (Lukasik et al., 2015; Niepoth et al., 2018) affect the probability that symbionts are lost from novel infections created in the laboratory. Aphids living on different species of host plant are likely to vary in their risk of exposure to natural enemies, and aphids in some environments might be more strongly selected to maintain their symbionts than others (Hrcek et al., 2018).

Overall, we find considerable variation in the density of symbionts in this system that is driven by both symbiont strain and host biotype. In addition, other abiotic and biotic factors that we did not explore here, such as the presence of other facultative endosymbiont species within a host (McLean et al., 2018), could also impact symbiont density. Our data suggest that the main relevance of density for this symbiosis is not through effects on the strength of symbiont mediated protection, but rather through its potential effects on symbiont transmission. Hosts will benefit from losing protective symbionts in environments where the risk of pathogen infection is low and keeping symbionts in other environments. Symbionts, in contrast, will be under pressure to avoid being lost from a host in all environments, but symbiont fitness is tied to that of its host, and trade-offs between density and host fitness might shape symbiont mechanisms that determine density. Together our data suggest that the optimal density of a *Regiella* infection is different from the perspective of aphid and symbiont fitness, and that the standing variation in density found in natural populations might be maintained in some systems by antagonistic evolutionary interactions between hosts and their microbes.

## Acknowledgements

We would like to thank Ciara Mann and Holly Nichols for valuable assistance with the experiments, Julia Ferrari for providing aphid genotypes, and Richard Humber and the USDA ARSEF collection of entomopathogenic fungi for providing the strains of *Pandora neoaphidis*. This work was funded by US NSF postdoctoral fellowship DBI-1306387 to BJP and UK NERC grant NE/K004972/1 to HCJG. BJP is a Pew Scholar in the Biomedical Sciences, supported by The Pew Charitable Trusts. JH was supported by Czech Science Foundation grant no. 17-27184Y.

## Statement of authorship

Designed research: BJP, JH, ACHM, HCJG. Performed research: BJP, JH, ACHM. Analyzed data: BJP, JAB. Wrote the paper: BJP, HCJG. Revised the manuscript critically for important intellectual content and approval of the version of the manuscript to be submitted: all authors.

## Figure and Table Legends

**Supplementary Figure 1:** Results of fungal infection using a second *Pandora* fungal genotype (ARSEF6631). Aphid genotypes are a subset of the lines used in the first infection. Error bars show standard error, and post-hoc tests comparing the interaction between genotype and symbiont status are shown along the top of the figure. Aphid biotypes are indicated by the colors of the genotype labels along the x-axis.

**Table S1:** Information on the collection location, year, symbiont background, and biotype of aphid lines used in this study.

| Host Genotype (lab code) | Location Collected | Year Collected | Original Symbionts | Biotype |
| --- | --- | --- | --- | --- |
| 502 | Maidens Green | 2003 | None | *Lathyrus pratensis* |
| 630 | Bernwood meadows | 2013 | None | *Lathyrus pratensis* |
| 663 | Bernwood meadows | 2014 | None | *Lotus corniculatus* |
| 664 | Bernwood meadows | 2014 | None | *Lotus corniculatus* |
| 11 | Whitby | 2012 | Ham | *Lotus pedunculatus* |
| 159 | Windsor | 2012 | Ham | *Lotus pedunculatus* |
| 74 | Windsor | 2012 | Ham | *Lotus pedunculatus* |
| 652 | Bourton-on-the-water | 2014 | None | *Medicago lupulina* |
| 672 | Chilswell (nr. Oxford) | 2014 | None | *Medicago lupulina* |
| 305 | Warburg Reserve | 2003 | Reg, Ham | *Medicago lupulina* |
| 207 | Lincoln | 2012 | Ham, X | *Medicago sativa* |
| 215 | Lincoln | 2012 | Reg | *Medicago sativa* |
| 302 | Lincoln | 2012 | Ham, X | *Medicago sativa* |
| 200 | Lincoln | 2012 | None | *Medicago sativa* |
| 222 | Whitby field | 2003 | Reg, Ham, Rick | *Medicago sativa* |
| 308 | Lincoln | 2012 | Ham | *Medicago sativa* |
| 123 | Wernershoehe | 2012 | Ham | *Ononis spinosa* |
| 256 | Speen | 2012 | Ham | *Ononis spinosa* |
| 257 | Speen | 2012 | Ham | *Ononis spinosa* |
| 133 | Cock Marsh | 2012 | Ham | *Ononis spinosa* |
| 607 | Wytham Wood | 2013 | Serr | *Pisum sativum* |
| 602 | Oxford Botanical Garden | 2014 | Serr | *Pisum sativum* |
| 622 | Barracks Lane, East Oxford | 2014 | None | *Pisum sativum* |
| 605 | Meadow Lane, East Oxford | 2013 | Serr | *Pisum sativum* |
| 619 | Barracks Lane, East Oxford | 2014 | Serr | *Pisum sativum* |
| 126 | Legoland | 2003 | Reg | *Trifolium spp.* |
| 319 | Oddington | 2012 | Reg | *Trifolium spp.* |
| 63 | Windsor Ranger's Gate | 2010 | Reg | *Trifolium spp.* |
| 317 | Oxfordshire | 2012 | Reg | *Trifolium spp.* |

**Table S2:** Results of post-hoc tests from the fungal infections. Statistically significant differences (P<0.05) are indicated in bold.

| Sporulation | Test Statistic | *p* |
| --- | --- | --- |
| *Lotus corniculatus* vs *Lotus pedunculatus* | $z = 1.090$ | 0.95471 |
| *Lotus corniculatus* vs *Lathyrus pratensis* | $z = 5.515$ | **< 0.001** |
| *Lotus corniculatus* vs *Medicago lupulina* | $z = 5.845$ | **< 0.001** |
| *Lotus corniculatus* vs *Medicago sativa* | $z = 0.624$ | 0.99839 |
| *Lotus corniculatus* vs *Ononis spinosa* | $z = 2.598$ | 0.14447 |
| *Lotus corniculatus* vs *Pisum sativum* | $z = -0.041$ | 1.00000 |
| *Lotus corniculatus* vs *Trifolium spp.* | $z = -0.610$ | 0.99860 |
| *Lotus pedunculatus* vs *Lathyrus pratensis* | $z = 5.249$ | **< 0.001** |
| *Lotus pedunculatus* vs *Medicago lupulina* | $z = 5.660$ | **< 0.001** |
| *Lotus pedunculatus* vs *Medicago sativa* | $z = -0.321$ | 0.99998 |
| *Lotus pedunculatus* vs *Ononis spinosa* | $z = 1.814$ | 0.58962 |
| *Lotus pedunculatus* vs *Pisum sativum* | $z = -0.847$ | 0.98919 |
| *Lotus pedunculatus* vs *Trifolium spp.* | $z = -1.240$ | 0.91211 |
| *Lathyrus pratensis* vs *Medicago lupulina* | $z = -0.071$ | 1.00000 |
| *Lathyrus pratensis* vs *Medicago sativa* | $z = -4.655$ | **< 0.001** |
| *Lathyrus pratensis* vs *Ononis spinosa* | $z = -4.043$ | **0.00120** |
| *Lathyrus pratensis* vs *Pisum sativum* | $z = -4.361$ | **< 0.001** |
| *Lathyrus pratensis* vs *Trifolium spp.* | $z = -3.960$ | **0.00165** |
| *Medicago lupulina* vs *Medicago sativa* | $z = -4.921$ | **< 0.001** |
| *Medicago lupulina* vs *Ononis spinosa* | $z = -4.404$ | **< 0.001** |
| *Medicago lupulina* vs *Pisum sativum* | $z = -4.538$ | **< 0.001** |
| *Medicago lupulina* vs *Trifolium spp.* | $z = -3.087$ | **0.03789** |
| *Medicago sativa* vs *Ononis spinosa* | $z = 1.700$ | 0.66743 |
| *Medicago sativa* vs *Pisum sativum* | $z = -0.535$ | 0.99940 |
| *Medicago sativa* vs *Trifolium spp.* | $z = -0.988$ | 0.97356 |
| *Ononis spinosa* vs *Pisum sativum* | $z = -1.919$ | 0.51651 |
| *Ononis spinosa* vs *Trifolium spp.* | $z = -2.036$ | 0.43541 |
| *Pisum sativum* vs *Trifolium spp.* | $z = -0.521$ | 0.99950 |
| Survival | Test Statistic | *p* |
| *Lotus corniculatus* vs *Lotus pedunculatus* | $z = 0.262$ | 1.00000 |
| *Lotus corniculatus* vs *Lathyrus pratensis* | $z = 1.502$ | 0.79900 |
| *Lotus corniculatus* vs *Medicago lupulina* | $z = 1.330$ | 0.88260 |
| *Lotus corniculatus* vs *Medicago sativa* | $z = -1.191$ | 0.93139 |
| *Lotus corniculatus* vs *Ononis spinosa* | $z = 0.223$ | 1.00000 |
| *Lotus corniculatus* vs *Pisum sativum* | $z = -1.813$ | 0.60083 |
| *Lotus corniculatus* vs *Trifolium spp.* | $z = 0.659$ | 0.99783 |
| *Lotus pedunculatus* vs *Lathyrus pratensis* | $z = 1.414$ | 0.84457 |
| *Lotus pedunculatus* vs *Medicago lupulina* | $z = 1.247$ | 0.91352 |
| *Lotus pedunculatus* vs *Medicago sativa* | $z = -1.716$ | 0.66633 |
| *Lotus pedunculatus* vs *Ononis spinosa* | $z = -0.067$ | 1.00000 |
| *Lotus pedunculatus* vs *Pisum sativum* | $z = -2.457$ | 0.20593 |
| *Lotus pedunculatus* vs *Trifolium spp.* | $z = 0.459$ | 0.99980 |
| *Lathyrus pratensis* vs *Medicago lupulina* | $z = -0.554$ | 0.99930 |
| *Lathyrus pratensis* vs *Medicago sativa* | $z = -2.683$ | 0.12220 |
| *Lathyrus pratensis* vs *Ononis spinosa* | $z = -1.527$ | 0.78465 |
| *Lathyrus pratensis* vs *Pisum sativum* | $z = -3.215$ | **0.02729** |
| *Lathyrus pratensis* vs *Trifolium spp.* | $z = -1.097$ | 0.95533 |
| *Medicago lupulina* vs *Medicago sativa* | z = -3.113 | **0.03691** |
| *Medicago lupulina* vs *Ononis spinosa* | z = -1.446 | 0.82921 |
| *Medicago lupulina* vs *Pisum sativum* | z = -3.908 | **0.00207** |
| *Medicago lupulina* vs *Trifolium spp.* | z = -1.425 | 0.83954 |
| *Medicago sativa* vs *Ononis spinosa* | z = 1.819 | 0.59546 |
| *Medicago sativa* vs *Pisum sativum* | z = -0.761 | 0.99463 |
| *Medicago sativa* vs *Trifolium spp.* | z = 2.238 | 0.31852 |
| *Ononis spinosa* vs *Pisum sativum* | z = -2.638 | 0.13613 |
| *Ononis spinosa* vs *Trifolium spp.* | z = 0.573 | 0.99912 |
| *Pisum sativum* vs *Trifolium spp.* | z = 3.003 | 0.05171 |

| Fungal infection with *Pandora* strain ARSEF6631 | Test Statistic | p |
| --- | --- | --- |
| Genotype 126 vs Genotype 200 | z = 0.043 | 1.00000 |
| Genotype 126 vs Genotype 619 | z = 0.042 | 1.00000 |
| Genotype 126 vs Genotype 74 | z = 0.042 | 1.00000 |
| Genotype 126 vs Genotype 652 | z = 0.049 | 1.00000 |
| Genotype 126 vs Genotype 305 | z = 0.048 | 1.00000 |
| Genotype 200 vs Genotype 619 | z = -0.381 | 0.99865 |
| Genotype 200 vs Genotype 74 | z = -0.048 | 1.00000 |
| Genotype 200 vs Genotype 652 | z = 5.320 | **< 0.001** |
| Genotype 200 vs Genotype 305 | z = 4.406 | **< 0.001** |
| Genotype 619 vs Genotype 74 | z = 0.381 | 0.99865 |
| Genotype 619 vs Genotype 652 | z = 6.434 | **< 0.001** |
| Genotype 619 vs Genotype 305 | z = 5.479 | **< 0.001** |
| Genotype 74 vs Genotype 652 | z = 5.237 | **< 0.001** |
| Genotype 74 vs Genotype 305 | z = 6.225 | **< 0.001** |
| Genotype 652 vs Genotype 305 | z = 1.384 | 0.69533 |

