## Supplementary figures and images for "Intraspecific variation in symbiont density in an insect-microbe symbiosis"

### Figure S1

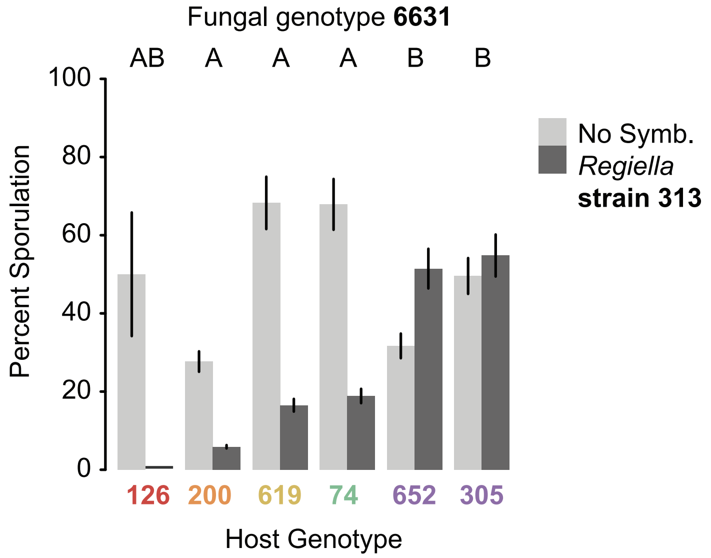
